# Microstructural characterization and validation of a 3D printed axon-mimetic phantom for diffusion MRI

**DOI:** 10.1101/2020.07.02.185397

**Authors:** Farah N. Mushtaha, Tristan K. Kuehn, Omar El-Deeb, Seyed A. Rohani, Luke W. Helpard, John Moore, Hanif Ladak, Amanda Moehring, Corey A. Baron, Ali R. Khan

## Abstract

**Purpose:** To introduce and characterize inexpensive and easily produced 3D-printed axon-mimetic (3AM) diffusion MRI (dMRI) phantoms in terms of pore geometry and diffusion kurtosis imaging (DKI) metrics.

**Methods:** Phantoms were 3D-printed with a composite printing material that, after dissolution of the PVA, exhibits microscopic fibrous pores. Confocal microscopy and synchrotron phase contrast micro-CT imaging were performed to visualize and assess the pore sizes. dMRI scans of four identical phantoms and phantoms with varying print parameters in water were performed at 9.4T. DKI was fit to both datasets and used to assess the reproducibility between phantoms and effects of print parameters on DKI metrics. Identical scans were performed 25 and 76 days later to test their stability.

**Results:** Segmentation of pores in three microscopy images yielded a mean, median, and standard deviation of equivalent pore diameters of 7.57 μm, 3.51 μm, and 12.13 μm, respectively. Phantoms with identical parameters showed a low coefficient of variation (∼10%) in DKI metrics (D=1.38 ×10^−3^ mm^2^/s and K=0.52, T1= 3960 ms and T2=119 ms). Printing temperature and speed had a small effect on DKI metrics (<16%) while infill density had a larger and more variable effect (>16%). The stability analysis showed small changes over 2.5 months (<7%).

**Conclusion:** 3AM phantoms can mimic the fibrous structure of axon bundles on a microscopic scale, serving as complex, anisotropic dMRI phantoms.

## 1. Introduction

Diffusion magnetic resonance imaging (dMRI) produces images by sensitizing the MRI signal to the random motion of water molecules on a micrometre scale. Models and representations of dMRI have been developed to describe the dMRI signal and produce metrics that are related to the microstructure and connectivity of the brain. To be clinically useful, the accuracy and reliability of dMRI-based model and representation parameters should be validated, but validation is difficult because there is typically no *in-vivo* ground truth for comparison.

A number of techniques have been used to produce dMRI ground truths, including numerical phantoms that simulate scan data (1,2), histology of brain samples scanned *ex-vivo* (3,4) or *in-vivo* before extraction (5–7), and physical phantoms with separately characterized microstructure (8,9). Numerical phantoms can use analytic models of diffusion in substrates composed of well-defined compartments (10), or Monte Carlo simulations of diffusion using arbitrarily complex and realistic mesh-based substrates (11–13). Numerical phantoms allow precise experimental control and increasingly realistic substrates, but realistic Monte Carlo simulations demand significant computational resources, which limits the possible volume of simulated substrates. Analysing histological sections of previously scanned brain tissue (14) provides data directly from a real brain, but histology is time-consuming and costly, covers a limited region of interest, and is difficult to register to a dMRI scan, especially when the scan was performed *in vivo*. Microstructural changes due to fixation and histological preparation limit the correspondence between these studies and *in-vivo* imaging. Physical phantoms are artificial objects designed to mimic the diffusion characteristics of the brain, ideally producing dMRI scan data similar to that seen from real brains. Physical phantoms occupy a middle ground between the two ends of a spectrum defined by numerical and *ex-vivo* studies: they produce real scan data with a well-known microstructural ground truth, but are not as customizable as numerical phantoms or as true to the structure of the brain as *ex-vivo* samples.

Several types of physical phantoms have been previously proposed. Glass capillaries (15) provide reliably anisotropic diffusion in a pattern that is straightforward to characterize, but cannot mimic some complex geometric fibre configurations that are observed in the brain. Plain (solid/non-hollow) fibres composed of a variety of materials are available on the market (16) with axon-scale diameters to mimic hindered diffusion between axons (17–19), but are difficult to arrange with the geometric complexity of some brain regions. Extruded or electrospun hollow fibers (20) mimic axonal diffusion patterns well, but require specialized equipment like high voltage power supplies (21) or melt-spinning extruders (22) to produce. As such, the development of dMRI phantoms involves trade-offs between the cost of materials and equipment, ease of production, the ability to achieve accurate brain-mimetic microstructure, and geometric complexity. Existing procedures to produce a physical phantom with biologically plausible microstructure and geometric complexity requires specialized equipment and/or is time-consuming, so an alternative that is both adequately brain-mimetic and easy to produce is needed.

We propose the use of Fused Deposition Modeling (FDM) 3D printing (23) as an inexpensive and flexible means of producing dMRI phantoms. To use FDM to produce 3D-printed Axon Mimetic (3AM) phantoms, we propose the use of a dual component “porous filament” material (24). In this work, we utilized “GEL-LAY” (LAY Filaments, Cologne, Germany), which is composed of a hydrophobic elastomeric matrix infused with pockets of polyvinyl alcohol (PVA). When 3D printed, the PVA forms long fibres within each line of printed composite material. PVA is water-soluble, so when this 3D printed material is immersed in water, the PVA fibres dissolve, leaving behind microscopic fibrous pores, as illustrated in Figure 1a. The anisotropic structure of these fibrous pores are similar to the anisotropic shape of axonal fibres and restrict diffusion in a similar way, potentially allowing 3D printed porous filaments to form the basis of a phantom that, due to the freedom to print along arbitrary directions, can potentially characterize the response of diffusion MRI representations (25) and models relative to fibre bending and crossing.

**Figure 1.**
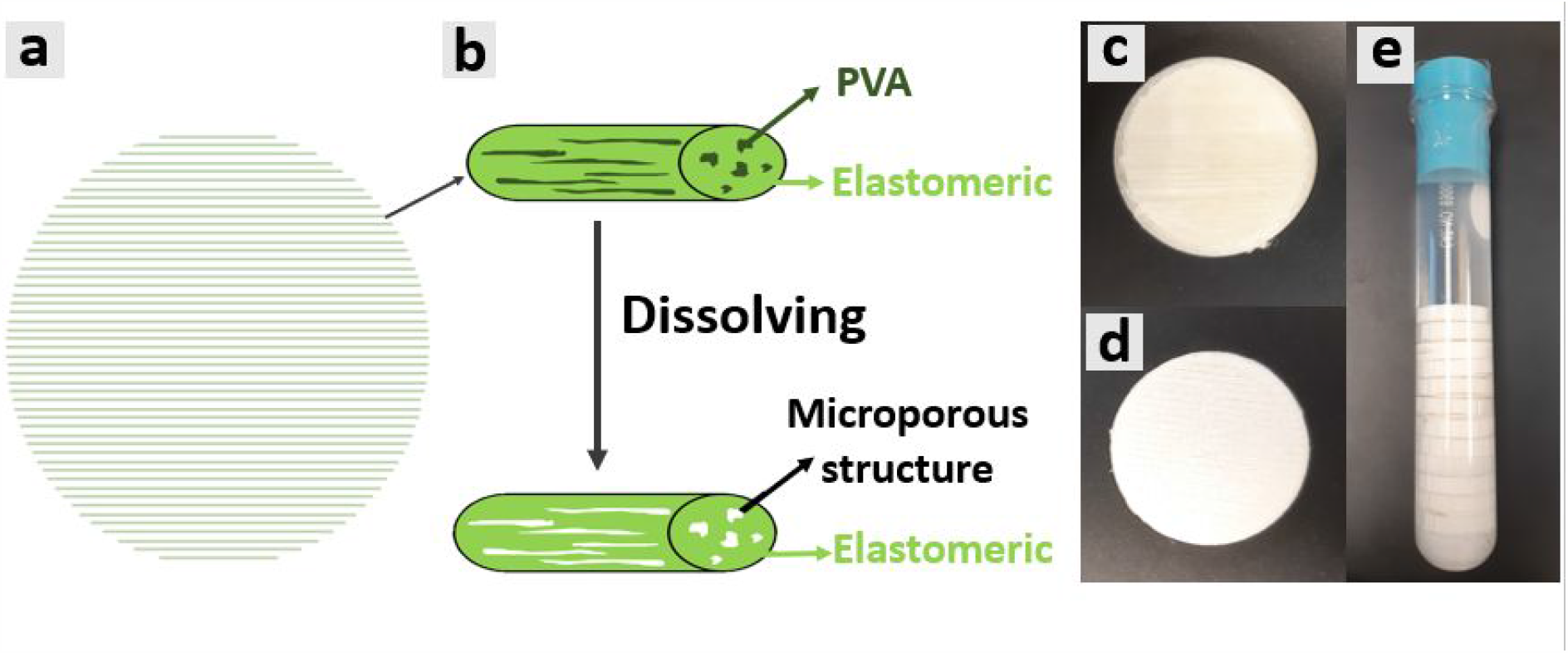
Illustration of 3AM phantom microstructure, and photos of printed phantoms. The cylinders in b) represent a single line of printed material. The disks in c-e) consist of hundreds of individual printed lines (as illustrated in a)) that are printed along the horizontal direction in a,c,d) and perpendicular to the long axis of the test-tube in e). a) 3AM Phantom printing schematic, where the material is printed along parallel lines. b) In each printed line, PVA dissolves away when placed in water leaving microporous structure. c) 3AM phantom before dissolving. d) 3AM phantom after dissolving. e) Dissolved 3AM phantoms stacked in a test tube with water that are ready for imaging.

In this study, we use both fluorescent microscopy and synchrotron phase-contrast micro-computed tomography, a novel technique that allows exceptionally fine-resolution 3D visualization of soft structures, to characterize and visualize the microstructure of 3AM phantoms. We examine the effects of FDM printing parameters on dMRI acquired in the phantoms, and assess the reproducibility and stability over time of 3AM phantoms’ dMRI characteristics. We also construct an analytic model of diffusion in 3AM phantoms based on the microscopy and micro-CT imaging and compare simulated diffusion signal from the analytic model to measured diffusion signal, assessing the accuracy of our characterization of 3AM phantoms’ microstructure.

## 2 Methods

### 2.1 Phantom preparation

An open protocol for producing 3AM phantoms has been developed and released as part of an Open Science Framework project (available in the Supporting information and hosted at osf.io/zrsp6). Required materials include an FDM 3D printer, a dual component “porous filament” material, a vacuum chamber, and a set of watertight containers. The protocol was used to produce all phantoms for this study.

All phantoms were designed using the open-source Ultimaker Cura software, and were printed with an Ultimaker 3 Extended 3D printer (Ultimaker, Geldermalsen, The Netherlands) loaded with GEL-LAY filament. Unless otherwise noted, phantoms were printed with printing parameters recommended by the material vendor (Table 1), and the same pattern of parallel lines of material in each layer.

**Table 1.** The parameters used to print the phantoms.

| Printing parameter | Low setting | Nominal setting | High setting |
| --- | --- | --- | --- |
| Printing temperature (°C) | 215 | 225 | 235 |
| Printing speed (mm/s) | 20 | 30 | 40 |
| Layer thickness (mm) | 0.06 | 0.1 | 0.14 |
| Infill density (%) | 75 | 100 | - |

After printing, phantoms to undergo dMRI scanning were immersed in 1L of room temperature water (∼23 °C) for 168 hours, then 20 mL of surfactant was added to decrease surface tension and allow the water to more easily enter the pores. The container was placed in a vacuum chamber at 1 bar for 48 hours to remove air bubbles. Finally, the phantoms were stacked in a test tube with distilled water for imaging (Figure 1b-d). Phantoms that undergo dMRI are kept in water at all times after preparation.

### 2.2 Microscopy

Three block-shaped 3AM phantoms with dimensions 25 mm × 15 mm × 35 mm were printed with vendor-recommended printing parameters. The blocks were immersed in water for 18 hours to soften by allowing a small portion of the PVA to dissolve, allowing them to reach the appropriate hardness level for slicing. Afterwards, the blocks were sliced into 50 μm slices in the plane transverse to the long axis of the pores using a Shandon Finesse ME+ microtome (ThermoScientific). The slices were immersed in deionized water for 3 hours to allow the remaining PVA to fully dissolve. They were then immersed in diluted Rhodamine Beta (Rhodamine B) and placed in a vacuum chamber at 1 bar for 20 minutes to infuse the dye into the slices and eliminate air bubbles. The slices were then mounted on positive microscope slides.

Confocal microscopy was performed to obtain high resolution images using a Leica SP5 laser system microscope with a 40X oil-immersion objective lens. Z-stacks of phantom slices were acquired using 1.0 μm step size for axial ranges between 5-17 μm. Using MATLAB’s image processing toolbox, three averaged z-stack confocal microscopy images were converted to grayscale (one from each block), then an adaptive thresholding technique (26) was used to compute a segmentation of each image. Finally, the equivalent radius of each region identified as a pore was calculated.

### 2.3 Synchrotron phase contrast micro-CT

A cylindrical phantom with a diameter of 5 mm and height of 4.5 mm was 3D printed, immersed in water for two days, and allowed to dry to improve CT contrast between the pores and the lattice. A propagation-based phase-contrast micro-computed tomography (micro-CT) scan was then performed in air at the Biomedical Imaging and Therapy beamline (05ID-2) at the Canadian Light Source Inc. (Saskatoon, Canada), using an energy of 30 keV, 3000 projections over 180°, 150 ms exposure time and 700 ms time per projection, total scan time of 1 hour, FOV=4096 × 4096 × 3456 μm^3^ (L × W × H), and an effective isotropic pixel size of 1.65 μm.

The micro-CT volume was segmented in two steps. First, the volume was rotated to align the lines of material with the axes of the volume, cropped to remove the edges of the sample, and downsampled to ⅛ of the initial resolution. The air-filled regions between lines of material were then segmented in the pre-processed volume by thresholding the standard deviation computed in local 3×3×3 windows. The resulting mask was then upsampled back to the source resolution.

To segment the pores in the micro-CT volume, an arbitrary x-z slice was chosen, and the air-filled region was masked out using the segmentation produced above. The resulting 2D image was segmented using an adaptive thresholding technique (26) from MATLAB’s image processing toolbox. The equivalent radius of each connected component identified as a pore was then calculated.

### 2.4 MRI scanning

Phantoms to undergo dMRI scanning were all scanned with the same parameters. Diffusion MRI was implemented with a 9.4 T Bruker small animal scanner using using a single-shot EPI sequence with 120 and 60 directions at b=2000 and 1000 s/mm^2^, respectively, 20 averages at b=0 s/mm^2^, diffusion gradient lobe duration (δ) of 4.062 ms, spacing between gradient lobes (Δ) of 13.062 ms, gradient magnitudes calculated to achieve the intended b-values, TE/TR=37/2500 ms, FOV=200×200 mm^2^, 0.7 mm isotropic in-plane resolution, and one 3 mm axial slice per phantom (8.5 min scan time).

For every resulting diffusion-weighted image, DiPy (27) was used to compute diffusion kurtosis imaging (DKI) metrics with a weighted ordinary least squares approach (28) at each voxel in an ROI manually drawn to avoid air bubbles in the phantoms.

A T1-mapping rapid acquisition relaxation enhancement (RARE) scan and a T2-mapping multi-slice-multi-echo (MSME) scan were performed on a set of eight phantoms with different 3D print parameters (Section 2.6). Both scans were implemented with a 9.4 T Bruker small animal scanner, FOV=40×40 mm^2^, 0.3125 mm isotropic in-plane resolution, and one 3 mm axial slice per phantom. Each slice was centered in a single phantom, mitigating the potential for partial volume effects between phantoms. The T2 mapping scan was performed with 16 echoes having TE ranging from 6.5 to 104 ms and TR=2000 ms. The T1 mapping scan was performed with TE = 5.7 ms, TRs of 500, 1000, 1500, 2000, and 3000 ms, and ETL = 2. To assess typical T1 and T2 values in 3AM phantoms, the mean and standard deviation T1 and T2 were calculated in the phantom produced with the nominal print parameters recommended by the manufacturer.

### 2.5 Phantoms for assessing the reproducibility of dMRI metrics

Four identical cylindrical 3AM phantoms were prepared with nominal printing parameters following the procedure outlined in Section 2.1. The phantoms were scanned with four axial slices (one slice per phantom), and DKI was fit to the scan data, all according to the imaging protocol outlined in Section 2.4. The coefficient of variation across the mean parameter values from the four phantoms was calculated for each DKI metric to assess the consistency of each metric across phantoms produced under identical conditions.

### 2.6 Phantoms for assessing stability over time and the effect of print parameters on dMRI

Eight cylindrical 3AM phantoms were produced according to the 3AM phantom protocol outlined in section 2.1, each with a width of 22 mm and height of 4.4 mm, composed of 44 layers of parallel lines.

Four FDM print parameters were altered across different phantoms to test their effects on the 3AM phantom’s microstructure. The four altered parameters include the temperature at which the phantoms are printed, the speed of printhead travel during the 3D print, the thickness of each layer printed in the phantom, and the infill density, which refers to the proportion of each layer taken up by material, and is altered by changing the distance between adjacent lines of material within each layer. One phantom was printed with the nominal print parameters recommended by the substrate manufacturer, and for each parameter, one phantom was printed with that parameter lower than nominal, and one phantom with that parameter higher than nominal (where applicable), as summarized in Table 1.

The eight phantoms were scanned with four axial slices and a scan time of 8.5 min per each of two dMRI scans to cover the test tube and DKI was fit to the data, all according to the protocol outlined in Section 2.5. The variation in each DKI metric was then assessed across the phantoms with each difference in print parameter.

To investigate the phantoms’ stability over longer time periods, identical dMRI scans and model fitting procedures were performed 25 and 76 days later, and the variations in each DKI metric in the phantom produced with nominal parameters across time was assessed.

### 2.7 Simulation

Camino (10,29,30) was used to construct an analytic diffusion model composed of two compartments: a free water compartment with a volume fraction estimated from the volume fraction of free water between lines of material in the micro-CT volume, and the second compartment composed of spins within parallel cylinders with radii distributed according to a gamma distribution. The parameters of the gamma distribution were determined using a maximum-likelihood fit of the equivalent pore diameters measured from the segmented microscopy images, which have a higher intrinsic resolution than the micro-CT. The simulation used a diffusivity of 2.1 ×10^−3^ mm^2^/s, the diffusivity of free water at room temperature. We simulated a diffusion MRI scan of our analytic model using the same experimental scan parameters we used for the phantoms and fit DKI to the simulated signal using DiPy, as we did for the experimental signal.

## 3. Results

### 3.1 Microscopy

Confocal microscopy images of slices in the plane transverse to the long axis of the pores revealed two types of pores present in the phantoms: larger pores (70-150 μm in diameter) that are believed to have been created by material tearing during the microscopy preparation process and gaps being left between lines of material (Figure 2), and smaller pores (1-30 μm in diameter) created by the dissolving of PVA fibres (Figure 3). The images also clearly showed the larger-scale arrangement of each line of material deposited during the 3D printed process.

**Figure 2.**
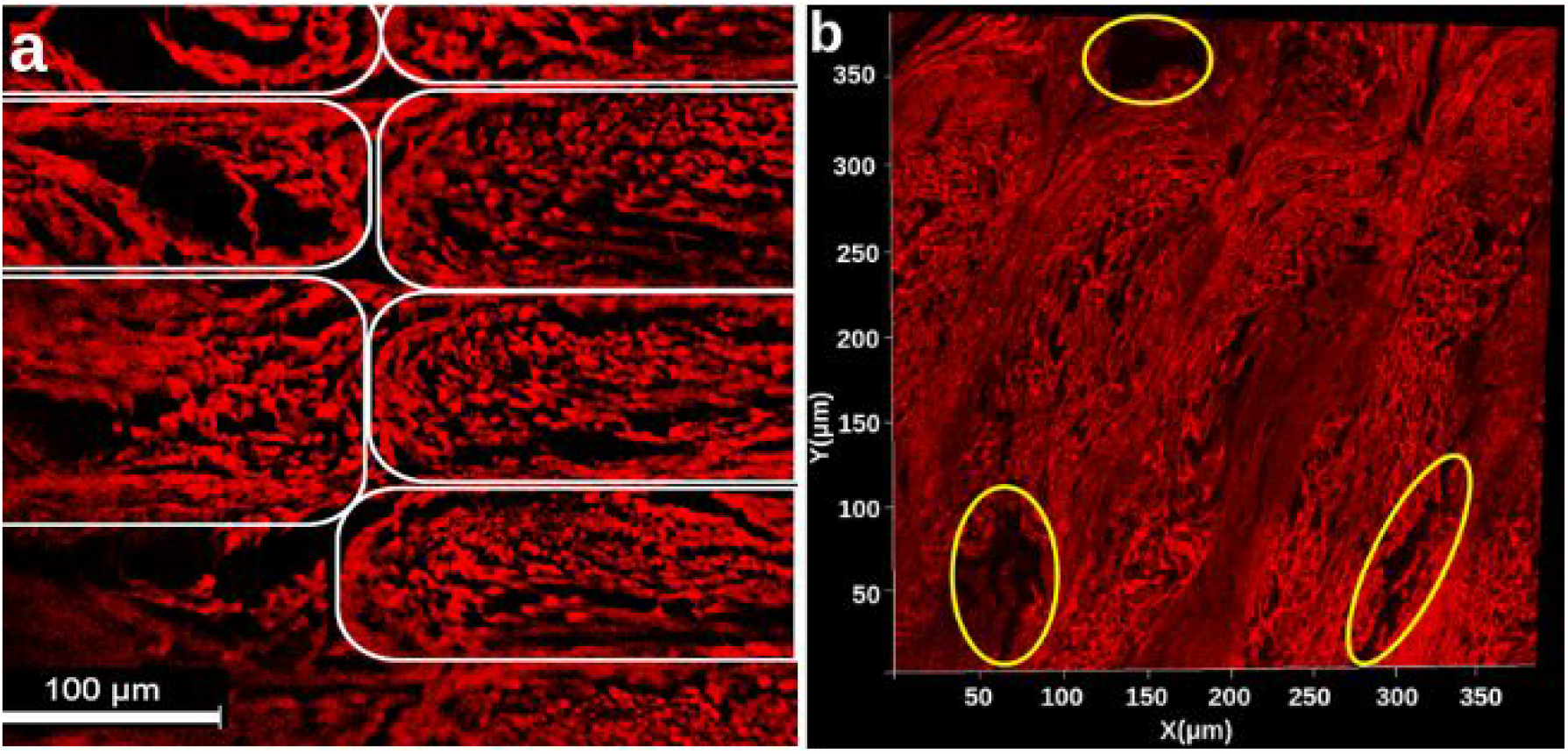
a) Confocal microscopy z-stack image of a stained cross sectional phantom sample, averaged across slices. Elastomeric matrix (red) and pores (black) are visible. Each white outline indicates an individual line of material, as depicted in Figure 1a. b) 2D projection of a 3D microscopy volume acquired with confocal microscopy. Regions shown in red are the matrix of the 3AM phantom that is composed of elastomer while the black regions are pores. Outlined in yellow are larger pores caused by the printing pattern of the phantom. In both a and b, the image plane is perpendicular to the long axis of the pores.

**Figure 3.**
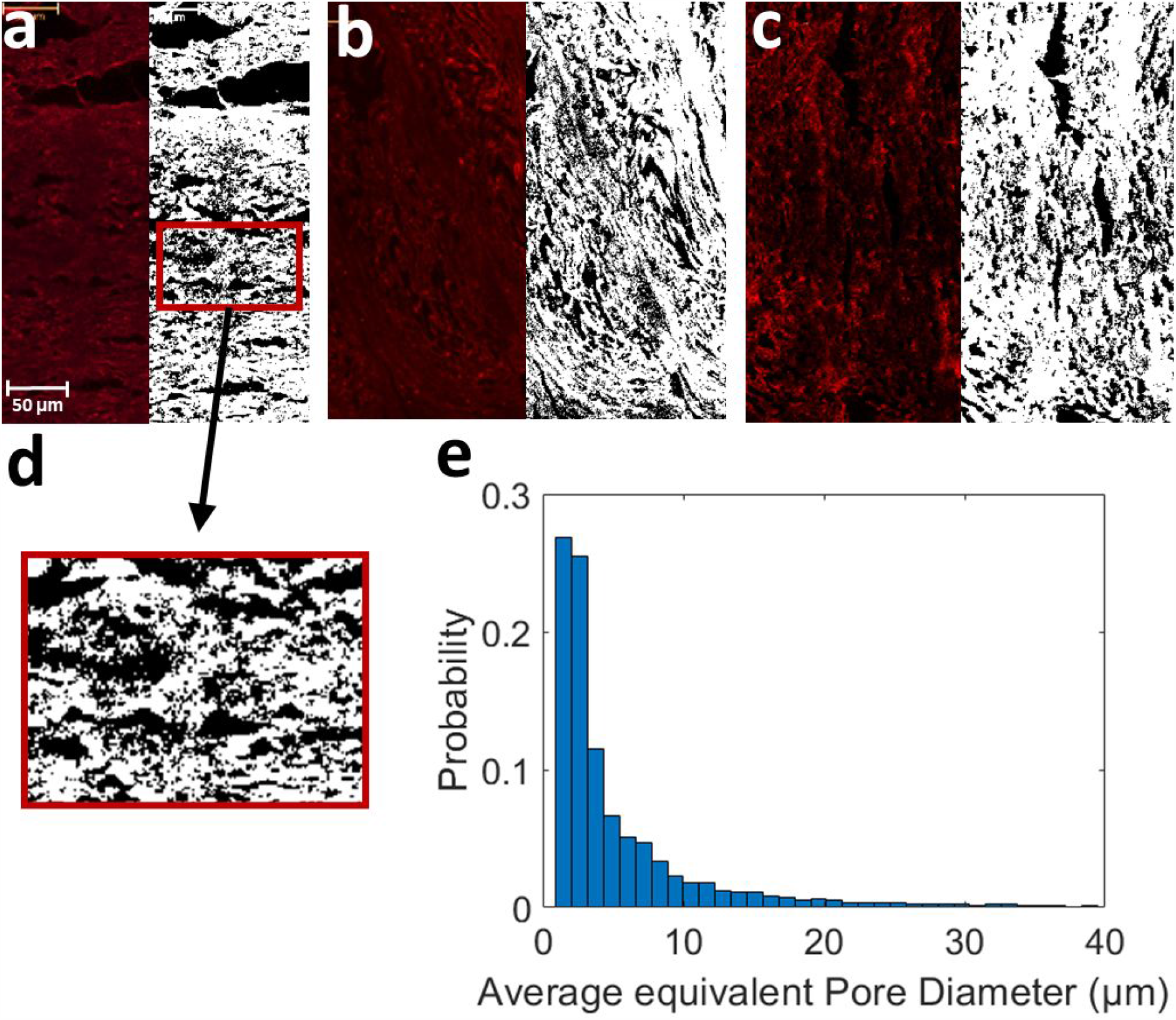
Pictures a-c, Confocal microscopy image before (left) and after (right) performing pore segmentation. Segmented pores are shown in black. d) zoomed in picture of segmented pores on a. e) Normalized histogram of pore equivalent diameters from the three segmentations in a-c (Total N = 10762).

The segmented pores from three averaged z-stack confocal microscopy images had a mean equivalent diameter of 7.57 μm, a median equivalent diameter of 3.51 μm, and a standard deviation of 12.13 μm (Figure 3).

### 3.2 Micro-CT

The propagation-based phase-contrast micro-CT image showed anisotropic pores that run parallel to the primary travel direction of the 3D print head (Figure 4), supporting the findings of the confocal microscopy. These pores had typical diameters on the order of ten microns and typical lengths in the hundreds of microns.

**Figure 4.**
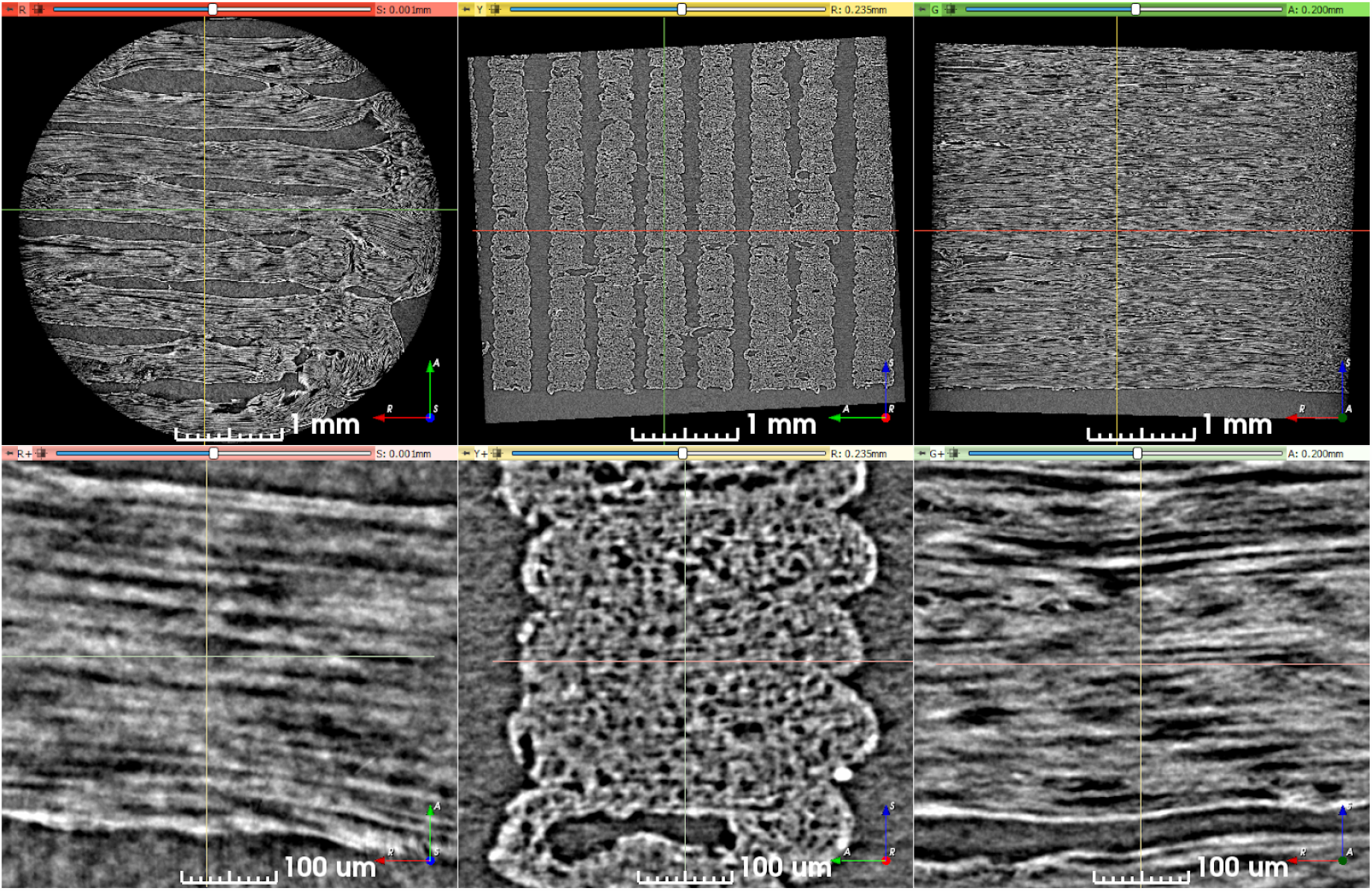
Synchrotron micro-CT scan data at two zoom levels, transformed to align lines of material with the viewing planes. Upper row: View of the entire scan ROI. Lower row: Detailed view of a short length of five stacked lines of material. The direction of print-head motion was left-right in the left- and right-most columns, and perpendicular to the image in the center column.

The segmented pores from one x-z slice of the micro-CT volume had a mean equivalent diameter of 8.69 μm, a median equivalent diameter of 5.71 μm, and a standard deviation of 20.40 μm (Figure 5).

**Figure 5.**
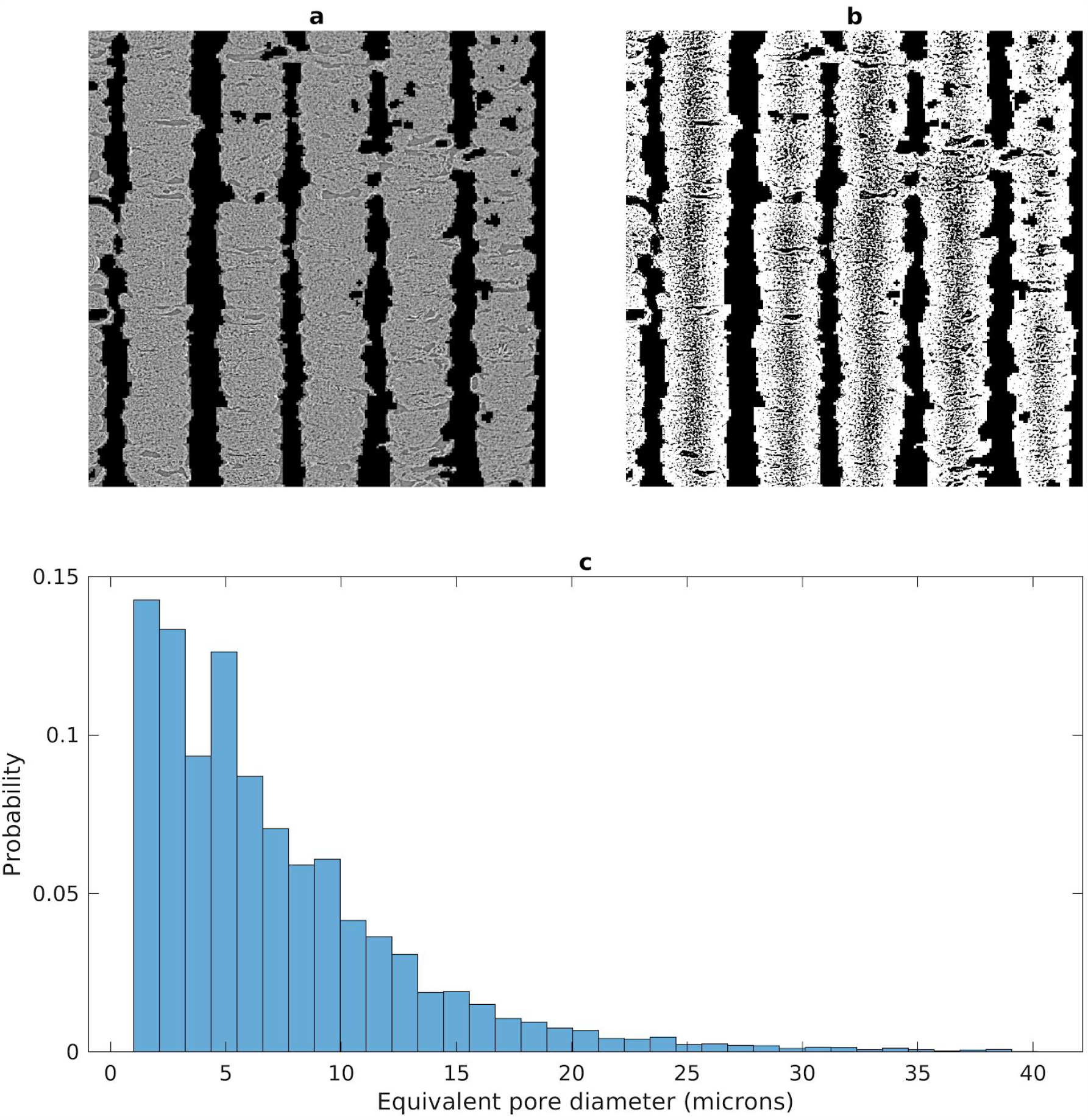
Segmentation of pores in a micro-CT slice. a) Preprocessed micro-CT slice with air-filled regions masked out. b) Segmentation results with pores shown in black. c) Histogram of pore equivalent diameters.

### 3.3 Nominal phantom characteristics and reproducibility

Examples of each image and metric map captured, with the ROI used, are shown in Figure 6. Some material leaked as the 3D print head traveled to reset between each printed layer, leaving a line of fibrous pores oriented nearly perpendicular to the intended pores. This line of leaked material is particularly prominent on the T2 and AK maps. This line only impacted a small number of voxels relative to the size of the mask, and likely had only a small effect on net measured parameters.

**Figure 6.**
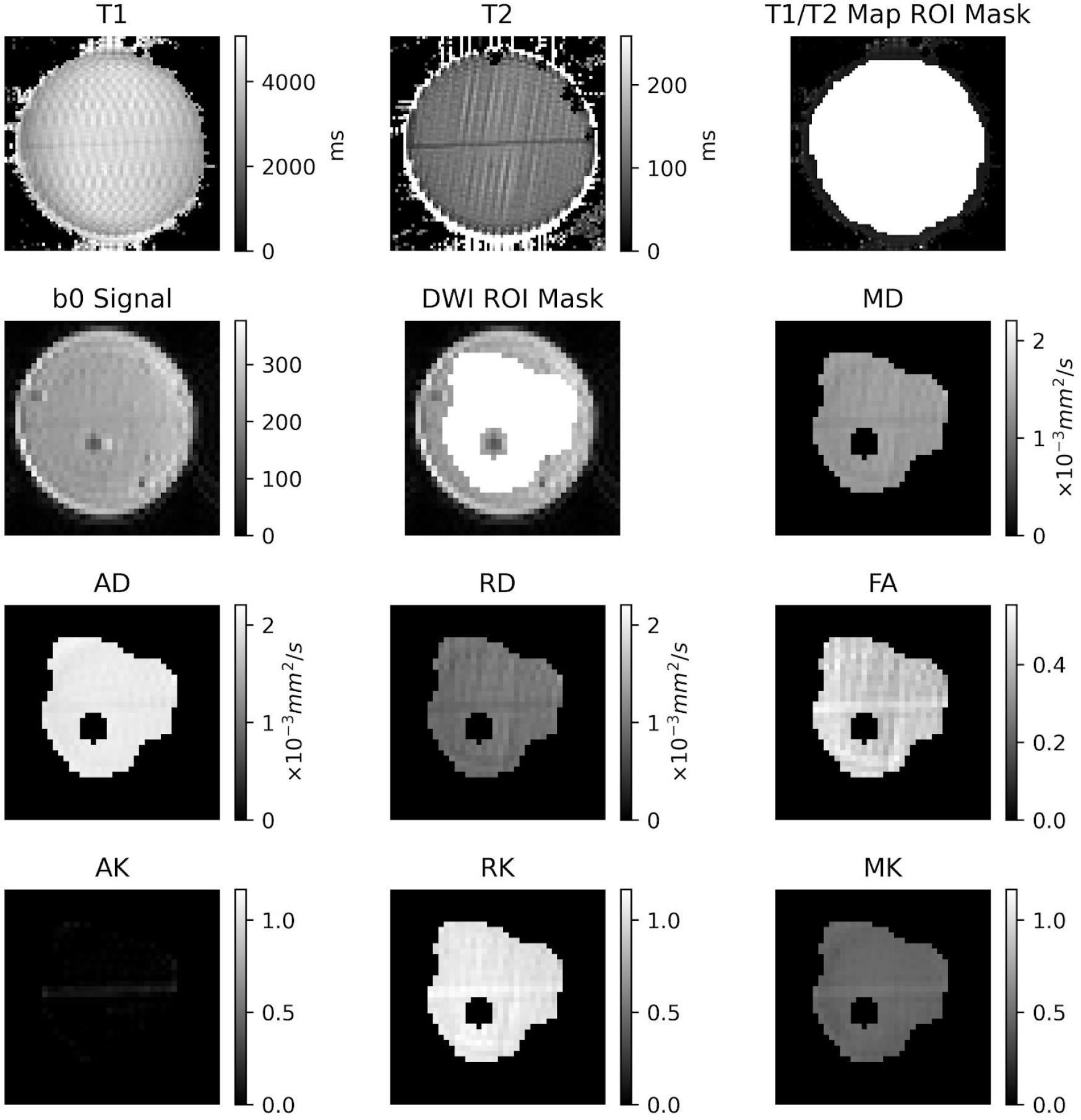
Examples of the produced maps in the nominal phantom. Top row: T1 and T2 maps, and ROI mask super imposed on the T1 map. Second row: Example non-diffusion weighted image, ROI mask superimposed on the non-diffusion weighted image, and mean diffusivity (MD). Third and fourth rows: DKI diffusivity metrics.

The mean T1 and T2 measured in a phantom printed with nominal printing parameters were 3960 ± 425 ms and 119 ± 21 ms, respectively.

The mean value of each DKI metric was calculated in each of the four nominal phantoms, then the mean and standard deviation for each DKI metric was calculated across those four values, as summarized in Table 2. The axial diffusivity (AD) is close to the diffusivity of pure water at room temperature (0.0022 mm^2^/s), and the radial diffusivity (RD) is about half the AD. The axial kurtosis (AK) is close to zero, suggesting little diffusion restriction in the direction parallel to the pores. The highest coefficient of variation across the four nominal phantoms was 15.00% for AK, and the rest were lower than 8%.

**Table 2.**
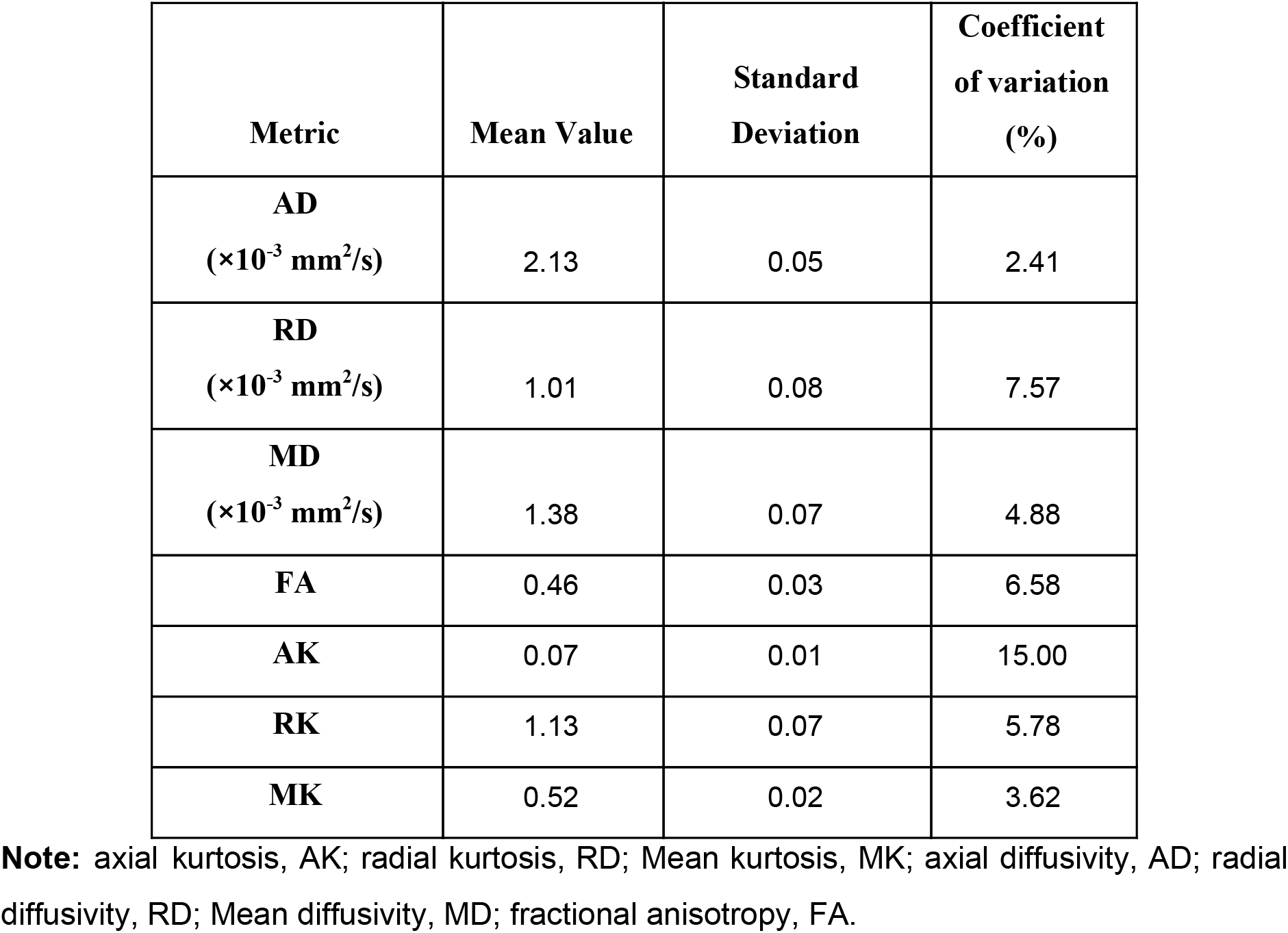
Mean dMRI metrics from 4 nominal phantoms and their coefficient of variation.

### 3.4 Metric variation with print parameters

Changes in the mean value of each metric with every print parameter were small for every print parameter except infill density (Figure 7). Excluding infill density, the difference from the nominal case was less than 16% for all metrics except AK, which had very high percent differences in some cases due to mean values close to zero. An increase in infill density of the phantom resulted in an increase in fractional anisotropy and a decrease in radial diffusivity.

**Figure 7.**
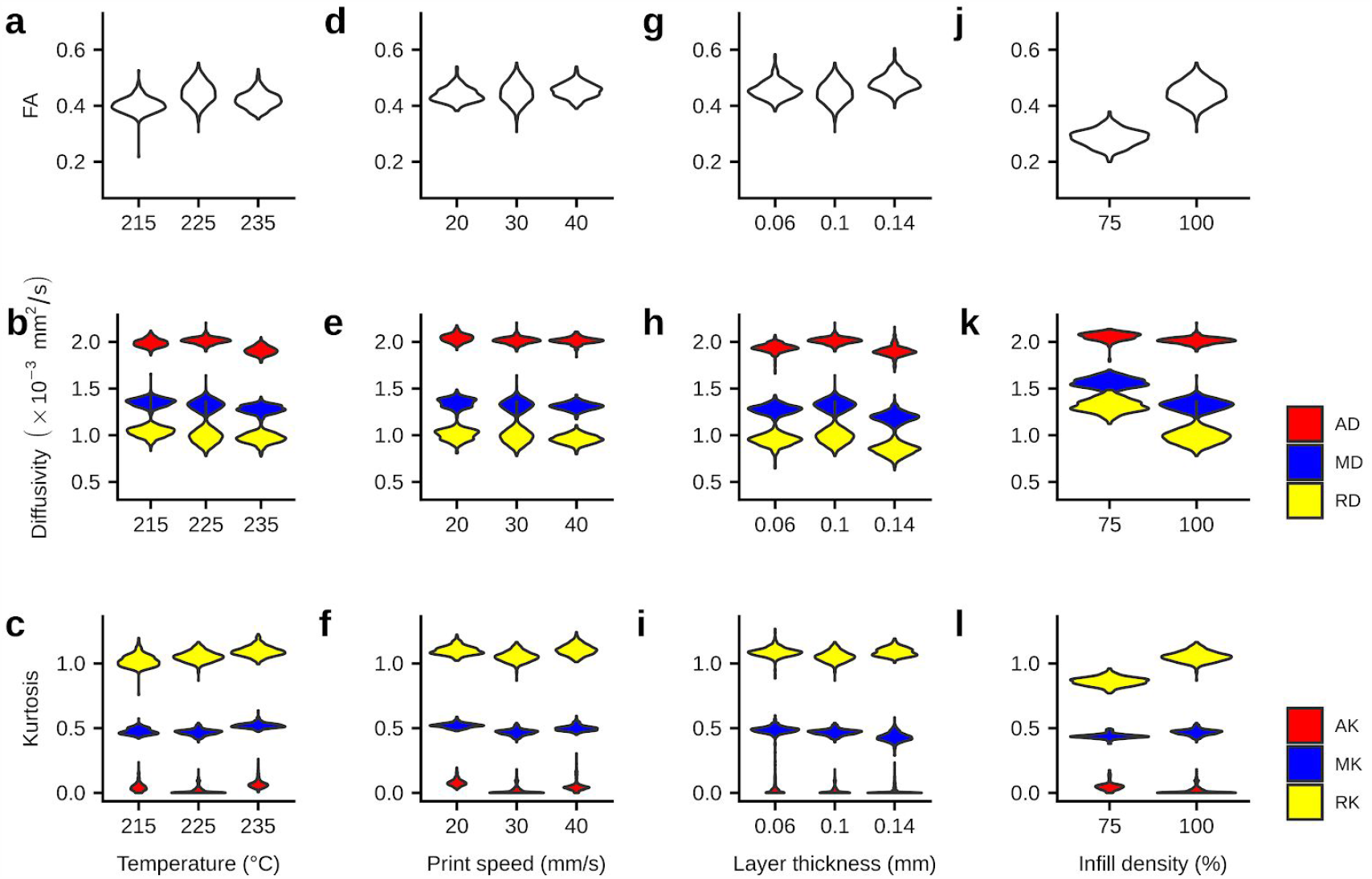
Mean FA, diffusivities, and kurtosis with different phantom printing parameters. a-c summarizes different metrics with different printing temperatures. d-f summarizes different metrics with different printing speeds. g-i summarizes different metrics with different layer thicknesses. j-l summarizes different metrics with different infill densities. Violin plots correspond to the distribution of a metric over all the voxels in a phantom.

### 3.5 Metric stability over time

Over the 76 days of study, only small differences in mean value were observed for all metrics (< 7%, except for AK due to some very small values of AK), as seen in Figure 8.

**Figure 8.**
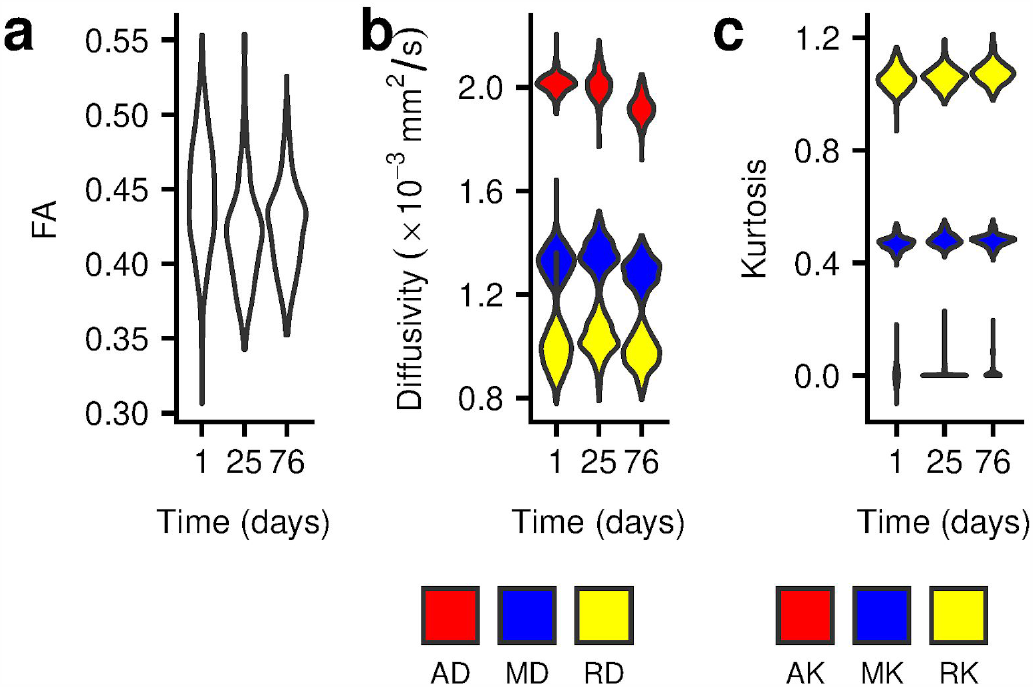
a) Mean Fractional anisotropy of a nominal phantom over 76 days. b) Mean Diffusivities of a nominal phantom over 76 days. c) Mean Kurtosis of a nominal phantom over 76 days. Error bars in all represent standard deviation of values in each corresponding phantom and scan time.

### 3.6 Simulation

From the micro-CT volume, we estimated a 34% volume fraction of free water in the phantom for the analytic model. The maximum likelihood fit of the gamma distribution to the array of equivalent pore diameters segmented from the microscopy images yielded the shape parameter a=1.18 and the scale parameter b=6.39 μm.

Fitting DKI to the simulated signal yielded a radial diffusivity of 0.944 ×10^−3^ mm^2^/s and radial kurtosis of 1.19, which is nearly identical to the experimentally observed values.

## 4. Discussion

### 4.1 Phantom vs. human microstructural fibre geometry

Microscopy images and synchrotron micro-CT images both indicate the presence of densely packed fibrous pores within a 3D printed composite material containing PVA within an elastomeric matrix. The close correspondence between simulated and measured diffusion behaviour suggests that the microscopy images resolve most of the phantoms’ pores, and that the pores are approximated well as straight cylinders.

A potential limitation of the pore size gamma distribution fitting is that primarily the tails of the histogram were used in the fitting. However, the smallest axons that were missed due to resolution limits only contribute to 0.11% of the total water volume, assuming the fitted gamma distribution parameters, and error likely had a small impact on diffusion parameters estimated from simulation. While the pores were generally not circular, the agreement of our simulations with experiment suggests that using an effective diameter based on the cross-sectional area may be an appropriate simplifying assumption. However, a notable limitation of these phantoms is that because they are composed of fibrous pores in a solid matrix, they have no analogue to the extra-axonal hindering of diffusion observed in the brain.

With a mean equivalent diameter of 7.56 μm and median equivalent diameter of 3.51 μm (Figure 3), these fibrous pores are larger than typical axons (31,32). This could have a considerable impact on using these phantoms to validate models that are sensitive to pore diameters (33); however, the impact would likely be small for long diffusion time acquisitions combined with models that assume “stick” diffusion (34,35). Nevertheless, the protocol introduced by this study produces long, straight pores that have a limited effect on diffusion along their path.

### 4.2 Phantom vs. human diffusion characteristics

The observed DKI metrics correspond well to those seen in real white matter but have some notable differences. The phantoms were scanned at room temperature, which has a lower diffusivity of free water (∼0.0022 mm^2^/s) than human body temperature (∼0.003 mm^2^/s) (36). This difference in the diffusivity of free water means that the diffusivity DKI metrics have a higher maximum possible value in vivo, which should be accounted for when analyzing those metrics.

AD in the phantoms was close to the diffusivity of free water at room temperature, while coherently organized white matter in the brain typically has AD of about 0.001 mm^2^/s, roughly a third of the diffusivity of free water at body temperature (37). This discrepancy indicates that the uniform fibre orientation and homogeneous microstructure of the phantoms results in a simpler axial diffusion environment than that observed in the brain. AK in the phantoms was close to zero, while practically all white matter regions in the brain have non-zero AK. This finding further supports the conclusion that there is an anatomically unrealistic homogeneity in the phantoms’ microstructure; real tissue is more heterogeneous and structurally complex than the 3AM phantoms.

RD in the 3AM phantoms was much lower than the diffusivity of free water at room temperature, indicating that diffusion is hindered and/or restricted perpendicular to the axis of the pores. The mean RK in the nominal 3AM phantom is non-zero but lower than typical values in orientationally coherent white matter tracts (38). This indicates that the fibrous pores of the 3AM phantom restrict diffusion but do not recreate the structural heterogeneity of real white matter. The increased pore diameters in 3AM phantoms compared to typical axon diameters mean that RD and RK will change with varying diffusion time; preliminary simulations suggest an RD decrease to 81% of the original value and RK increase to 135% of the original value for a doubling of diffusion time from 13 ms to 26 ms.

MD in the 3AM phantoms was about 63% of the diffusivity of free water at room temperature, while MD in real white matter is typically less than 33% of the diffusivity of water at body temperature. Partially due to the near-total lack of restriction or hindering of diffusion in the axial direction, there is less restriction/hindering of diffusion overall in 3AM phantoms than in white matter. MK in the 3AM phantoms was about half the typical values in human white matter (38), further indicating that 3AM phantoms have less heterogeneous microstructure than real axonal tracts.

The FA is within the range of typical values observed in the human white matter (37), but lower than that observed in the most coherent regions like the corpus callosum (39). The micro-CT images show gaps between adjacent lines of material even at 100% infill density, which leads to a non-negligible free water compartment within the phantoms of 34%. This free water compartment reduces the overall FA due to partial voluming, which suggests that a higher FA may be achievable with an altered 3D printing process. This free-water compartment likely also contributes to the relatively low kurtosis values and the diffusivities being relatively close to the free water value.

### 4.3 Phantom reproducibility and stability

Infill density was the only print parameter that had a large effect on any of the DKI metrics, with its greatest effect on FA, RD, and RK. The likely explanation for this effect is that a lower infill density replaces the elastomeric matrix with free water, increasing the effect of partial voluming between the two components.

The small effect of the other 3D print parameters on the observed DKI metrics suggests that minor variations in print parameters across different 3D printers should not greatly affect the characteristics of the phantoms they print. However, this also means that it is likely not possible to change print parameters to tune a phantom’s diffusion characteristics. That said, it may be possible to tailor diffusion characteristics using different porous filament materials. For example, the PORO-LAY filament line contains several filament types that have a porous microstructure that is created by PVA dissolving away, but they differ in the composition of the elastomer, including its hardness level. The low coefficient of variation across identical phantoms supports the conclusion that the 3AM phantom production protocol is repeatable, at least across multiple prints on a single printer with the same material.

The stability analysis showed only small changes over the time period of 2.5 months. While the scanner is located in a temperature controlled facility and it is unlikely that there were substantial temperature changes, it is possible that variation in these temperatures could explain the downward trend in AD over time. Nevertheless, the demonstrated stability of the phantom microstructure over a relatively long time period suggests that one prepared phantom sample can at least be transported to multiple sites in multi-centre studies without concern for parameter changes between scans.

### 4.4 Method advantages

The phantoms we have introduced are produced with FDM 3D printing, a widely accessible and inexpensive production technique. This potentially allows customizability of phantoms microstructural directionality to create crossing fibre bundles in multiple directions. A single spool of printing material (∼ $50 USD) can be used to produce hundreds of phantoms. The development of this technique lowers the barrier to entry for researchers to conduct phantom studies for validation of certain dMRI models of white matter. Such studies bridge the gap between simulations and studies using fixed tissue, serving as a useful option for the dMRI modeling community.

Compared to existing phantoms, the primary advantage of 3AM phantoms is that they can be manufactured without specialized equipment. 3AM complement plain fibre phantoms by providing a diffusion environment analogous to intra-axonal diffusion. Furthermore, 3AM fibres do not require external frames or molds to manipulate the arrangement of their fibrous pores, like plain fibre phantoms or extruded hollow fibre phantoms do. This ease of production comes with the limitation that the pore diameter distribution is not customizable, and that it produces pores with a larger diameter than real axons.

The analyses performed for this study show the existence of axon-mimetic fibrous pores in 3AM phantoms that modulate dMRI signal to approximate white matter anatomy. By analyzing the microstructure of the phantoms with both confocal microscopy and phase contrast micro-CT, we have confirmed that the fibrous pores in our phantoms are of an appropriate size and shape to mimic axonal fibres. These findings are supported by the observation of non-zero FA and RK. Furthermore, the quantitative T_1_ and T_2_ scans indicate that the phantoms do not shorten relaxation times enough to significantly reduce SNR in dMRI scans, and they can be reduced via doping (e.g., copper II sulphate) to better agree with values found in tissue.

FDM offers the flexibility to alter the print-head direction between layers, which may enable the exploration of crossing fibre effects and their associated models. In our preliminary investigations, FA reductions are observed with increasing fibre crossing angle. Future work will investigate optimal phantom design for validation of crossing fibre models.

## 5 Conclusion

In this study we introduce 3AM phantoms, a novel class of dMRI phantoms that are affordable and straight-forward to produce. 3AM phantoms mimic the microporous structure of axon bundles in white matter, and the utilization of 3D printing likely opens the door for customizability and the production of complex, anatomically accurate phantoms.

## Acknowledgements

This work was supported by the Canada First Research Excellence Fund, Brain Canada, and Discovery Grants from the Natural Sciences and Engineering Research Council (NSERC). Part of the research described in this paper was conducted at the BioMedical Imaging and Therapy (BMIT) facility at the Canadian Light Source Inc. (CLSI), which is funded by the Canada Foundation for Innovation, the Natural Sciences and Engineering Research Council of Canada, the National Research Council Canada, the Canadian Institutes of Health Research, the Government of Saskatchewan, Western Economic Diversification Canada, and the University of Saskatchewan.

